# Sparse Infomax Based on Hoyer Projection and its Application to Simulated Structural MRI and SNP Data

**DOI:** 10.1101/570317

**Authors:** Kuaikuai Duan, Rogers F. Silva, Jiayu Chen, Dongdong Lin, Vince D. Calhoun, Jingyu Liu

**Affiliations:** Department of Electrical and Computer Engineering, The University of New Mexico, USA; The Mind Research Network and Lovelace Biomedical and Environmental Research Institute, USA

**Keywords:** Sparse infomax, Hoyer projection, Imaging data, SNP data, Pattern recovery

## Abstract

Independent component analysis has been widely applied to brain imaging and genetic data analyses for its ability to identify interpretable latent sources. Nevertheless, leveraging source sparsity in a more granular way may further improve its ability to optimize the solution for certain data types. For this purpose, we propose a sparse infomax algorithm based on nonlinear Hoyer projection, leveraging both sparsity and statistical independence of latent sources. The proposed algorithm iteratively updates the unmixing matrix by infomax (for independence) and the sources by Hoyer projection (for sparsity), feeding the sparse sources back as input data for the next iteration. Consequently, sparseness propagates effectively through infomax iterations, producing sources with more desirable properties. Simulation results on both brain imaging and genetic data demonstrate that the proposed algorithm yields improved pattern recovery, particularly under low signal-to-noise ratio conditions, as well as improved sparseness compared to traditional infomax.

## 1. INTRODUCTION

Blind source separation methods, such as independent component analysis (ICA), have been widely applied to brain magnetic resonance imaging (MRI) and electroencephalogram data, as well as genetic data, etc. Infomax, as one of the most robust ICA approaches, has shown great ability to recover super-Gaussian sources when the signal-to-noise (or - background) ratio (SNR or SBR) of data is high[1]. Meanwhile, improvement is needed for the situation where the SNR or SBR is low, such as genetic mutation data, namely single nucleotide polymorphisms (SNPs). SNPs are categorical values (0, 1, or 2) that indicate the number of minor (usually mutated) alleles at each SNP locus. Typically, the variance of SNP independent sources of interest (modeling genetic interactions) is similar to that of the background (presenting genetic coding for unknown or less represented biological processes). In such case, other source properties could be leveraged to improve the ICA performance.

Sparsity is one such property that is commonly incorporated into ICA approaches. Ge et. al [2] proposed sparse FastICA based on the smoothed *l*_0_ norm, which outperformed FastICA in terms of source detection and robustness to noise. However, its performance was still similar to that of infomax, likely because infomax already includes an implicit (non-tunable) sparsity prior due to the choice of nonlinear neuronal activation function (sigmoid or tanh), which optimizes for super-Gaussian distributions.

In order to add tunable sparsity control to infomax, we consider the Hoyer index. Hurley and Rickard [3] have shown that the Gini index is the only one satisfying the six axiomatic attributes of sparsity measures. But optimizing the Gini index is difficult due to its built-in sorting operation. The Hoyer index is a sparsity measure second only to the Gini index, and easy to optimize. Compared to standard *l*_1_ or *l*_2_ norms, the Hoyer index is an extension of the ratio between *l*_1_ and *l*_2_. Also, it is scale invariant and has no need for additional normalization, which presents great potential in the context of ICA algorithms as the scale of source estimates is arbitrary and irrelevant for the pursuit of independence.

In this paper, we propose a sparse infomax algorithm based on the nonlinear Hoyer projection. Sparse infomax updates the unmixing matrix the same way as infomax optimization, but applies the Hoyer projection to the source estimates before propagating them as mixed data for the next iteration.

## 2. METHODS

For a vector **z**, its Hoyer index is defined as follows:

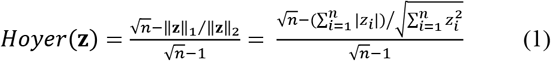

where *n* is the number of elements in **z**. The Hoyer index ranges between 0 and 1. A Hoyer index of 0 (least sparse) indicates that all elements in **z** are the same, while a Hoyer index of 1 (most sparse) means that there is only one non-zero element in **z**.

The proposed sparse infomax has the same cost function as traditional infomax[1] (i.e. maximizing the differential entropy H(**y**) with respect to the unmixing weight matrix **W**) except for the additional constraint on the estimated sources **S**, calling for a *Hoyer*(**s**) larger or equal than the preset threshold *t*; see (2). Assume that the observed data **X** is a linear mixture of independent sources (**X** = **AS**), where **A** is the mixing matrix. Then the source **S** can be estimated as **WX**, where **W** = **A**^−1^. Since the Hoyer index[4] constraint is optimized through a series of nonlinear projections, **A** cannot be estimated directly by **W**^−1^ after convergence. Instead, Tychonov-regularized least squares [5] is employed to estimate **A** using the sources estimated by sparse infomax. Tychonov-regularized least squares takes the data noise into account, providing a better solution than traditional least squares. Its cost function is defined in (3), where *δ* is the regularization parameter, **a** is a row of **A**, and **x**^T^ represents a row of **X**. The overall mathematical model of sparse infomax is defined below:

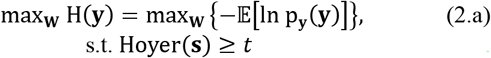

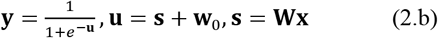

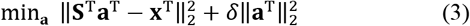

where p_**y**_(**y**) is the probability density function of the output vector **y** (the nonlinear transformation of the bias-adjusted source **u**), and **X** = **AS**, **S** = **WX**. Infomax is solved by stochastic gradient descent (SGD), using the natural gradient of (2.a) with respect to **W** and the traditional gradient of (2.a) with respect to **w**_**0**_:

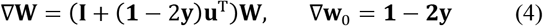

where **I** is an identity matrix and **1** is a vector of ones. T is the transpose operation. The pseudocode for the proposed sparse infomax algorithm is described as follows:

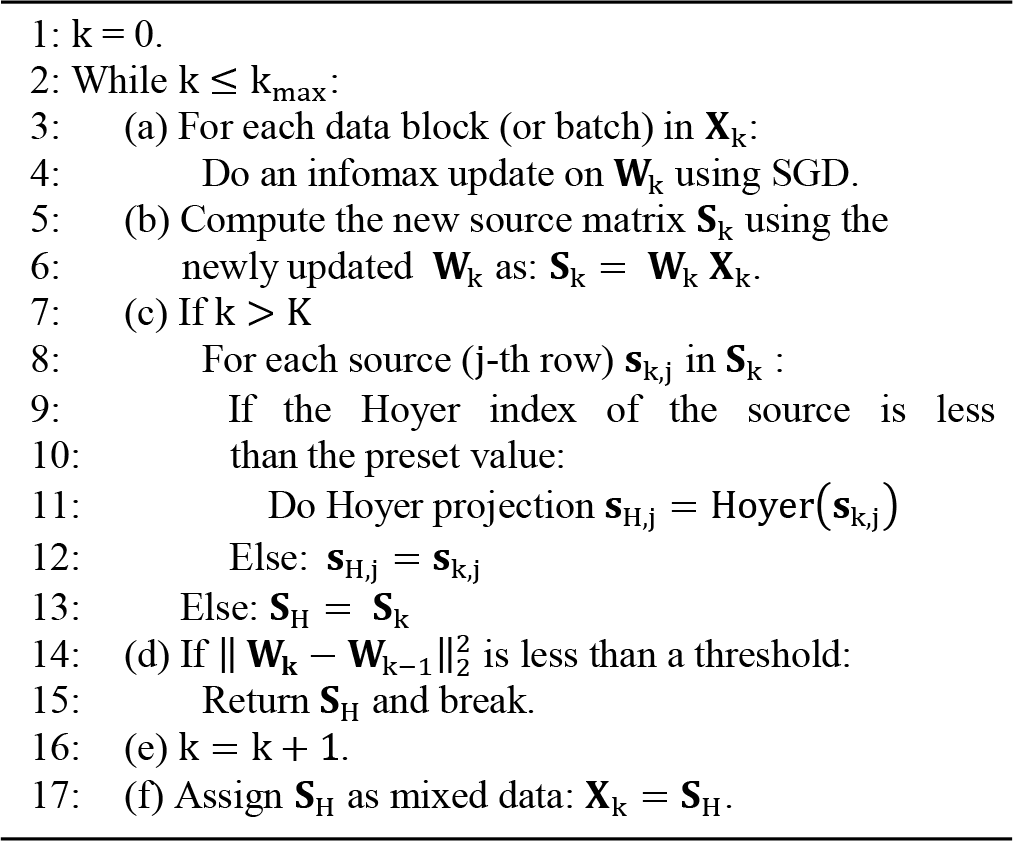

In the proposed sparse infomax, the independence and sparsity of sources are optimized iteratively. First, the unmixing matrix **W** is updated based on infomax’s natural gradient. For the first K steps (default is K = 5), only infomax update is applied to prioritize independence at the initial stage. For the ensuing iterations, after updating the **W** matrix, the source matrix **S** = **WX** is estimated. Then, we compute the Hoyer index for each source and perform Hoyer projection if the source sparsity does not meet the preset value for Hoyer index. The Hoyer projection is implemented based on Hoyer’s paper [4], which keeps the *l*_2_ norm of sources unchanged while updating the elements of each source (a row of **S**) until the *l*_1_ norm reaches a value that yields the desired sparseness. Contrary to the underlying linear generative model of infomax, the Hoyer projection is a nonlinear transformation; the update on the elements of each source involves zeroing out small values and increasing large ones. After Hoyer projection, we propagate the updated sources as mixed data for the next iteration. During Hoyer projection, we set the desired Hoyer index as the current Hoyer value of the source plus 0.05, if it is not greater than the preset value. Otherwise, we employ the preset value as the each subject). desired Hoyer index. This is to avoid abrupt alterations of the sources.

The solution to the Tychonov-regularized least squares is found by solving the following linear equation:

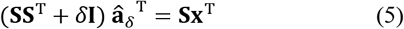

where 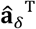 is the estimated transpose of **a**. The role of *δ***I** is to stabilize the least squares solution for noisy data. As suggested by Strang in [5], the value of *δ* can be chosen as 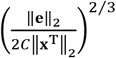 (and the constant *C* can be approximated by 1/*σ*^3^, where *σ* is the smallest singular value of **S**. In sparse infomax, we evaluate *C* based on the final estimated source matrix **S**. The noise *e* can be approximated using the background of the observed mixed data **x**^T^ after excluding elements in the signal region based on the estimated sources. Finally, we plug *δ* into (5) to obtain the estimated 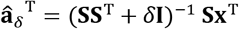 for each row of **A** (here, for each subject).

## 3. SIMULATIONS AND RESULTS

### 3.1 Structural MRI

The SimTB Toolbox (http://mialab.mrn.org/software/simtb/) [6] was used to simulate seven non-overlapping structural MRI (sMRI) components, each with 31064 voxels and similar sparsity levels (Hoyer indices were all around 0.85). By default, SimTB normalized the component values to be between 0 and 1. As components generated by SimTB had continuous intensity values within the whole brain, we set voxels with intensity levels less than 0.25 as 0 to refine the components, yielding the source matrix **S**_7×31064_ (plotted in Fig. 1), and only regions with intensity values larger than 0.25 were treated as the ground-truth component regions (GTCR).

**Fig. 1.**
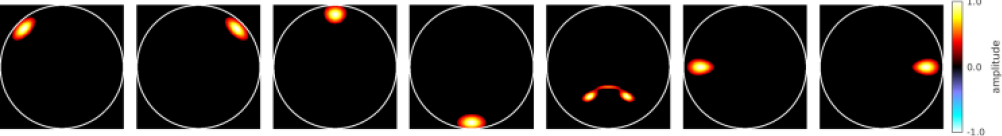
Seven non-overlapping sMRI components.

The mixing matrix **A**_200×7_ was randomly generated from a uniform distribution 𝒰(0,1) to represent the components expression levels for 200 subjects. Then, we obtained the noiseless data matrix as **X**_*clean*_ = **AS**. Rician noise with a contrast-to-noise ratio (CNR, as defined in [6]) of 3 was added to **X**_*clean*_ in order to obtain the final mixed data **X**.

Subject-wise mean removal and principal component analysis (PCA) were applied to data **X** and seven principal components were extracted for further infomax or sparse infomax analyses. To test the performance of sparse infomax, we applied it with the preset Hoyer index value varying from 0.4 to 0.95, with a step size of 0.05. Various component-wise performance measures were evaluated, including correlations between the estimated and ground-truth components, and correlations between the estimated and ground-truth mixing matrix, sensitivity, specificity and F1 scores, sparsity measures (Hoyer/Gini indices), as well as SBR. In Fig. 2, we reported average values across components. Here, SBR was defined as:

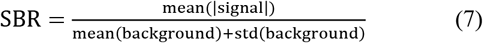

where std denotes the standard deviation. Signal and background regions were defined based on both GTCR and a component-level z-threshold of |*Z*| = 3.5 (same for SNP data analysis). Thus, assuming that the component background region follows a normal distribution, SBR measures how far on average a component’s signal is away from the background. Higher SBR values indicate better signal discrimination.

**Fig. 2.**
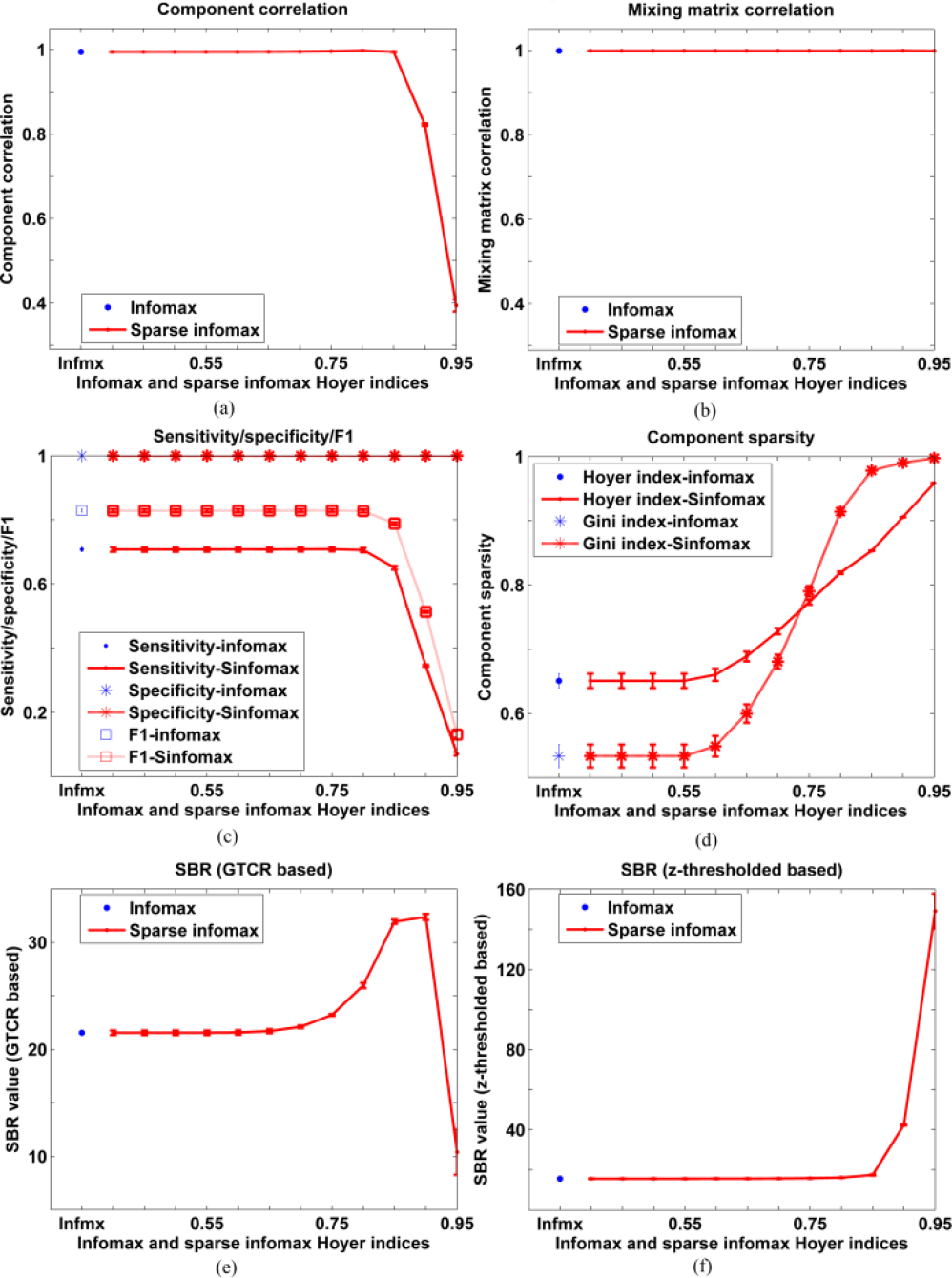
Infomax and sparse infomax performances on simulated sMRI data (a) component correlation, (b) mixing matrix correlation, (c) sensitivity/specificity/F1 score, (d) Gini/Hoyer indices, (e) SBR (GTCR based), (f) SBR (z-threshold based, |*z*| > 3.5). Notes: Infmx denotes infomax, and 0.4 to 0.95 are the preset Hoyer index values used in sparse infomax; Sinfomax in (c) and (d) denotes sparse infomax (the same for Fig.3, Fig.4 and Fig.5). The error bars reflect mean ± standard error across seven components (the same for Fig.3, Fig.4 and Fig.5).

Fig. 2 shows that, except for sparseness (Gini/Hoyer indices) all performance measures are comparable between infomax and sparse infomax when the preset Hoyer index value is reasonable (here, less than the true value of 0.85). In all cases, the sparsity measures of components from sparse infomax are improved compared to those from infomax. Furthermore, in the very low SNR cases (e.g. CNR=0.5), we also observe improvement in component correlation and SBR by sparse infomax, which are in agreement with the following SNP data results. Thus, they are omitted here. The convergence speed (iterations taken until convergence) is 80 iterations for infomax, and 71 for sparse infomax (mean of all Hoyer indices scenarios).

### 3.2 SNP data

We simulated SNP data of 10000 loci with 7 non-overlapping SNP component patterns using Plink 1.9 (http://zzz.bwh.harvard.edu/plink/) for three scenarios:

#### Scenario 1

for each SNP component, the effect size = 3.5, number of risk SNP = 150 (signal region), and sample size (number of subjects) varying from 100 to 1000 (100, 200, 300, 500, 700, 1000). Effect size (ES) is defined as follow:

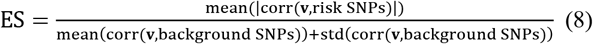

where **v** is the representative risk SNP, corr and std denote correlation and standard deviation operations, respectively. Subject-wise mean removal and PCA were applied to SNP data to generate 8 principal components, including one extra for noise/background. Subsequently, infomax and sparse infomax with preset Hoyer value of 0.4 were performed to estimate 8 independent components. Performance measures including correlation between the estimated and ground-truth components, sensitivity, specificity and F1 scores, sparsity measures (Hoyer/Gini indices), as well as SBR (z-threshold of |*Z*| > 3.5), were computed and averaged across components for both methods (Fig. 3).

**Fig. 3.**
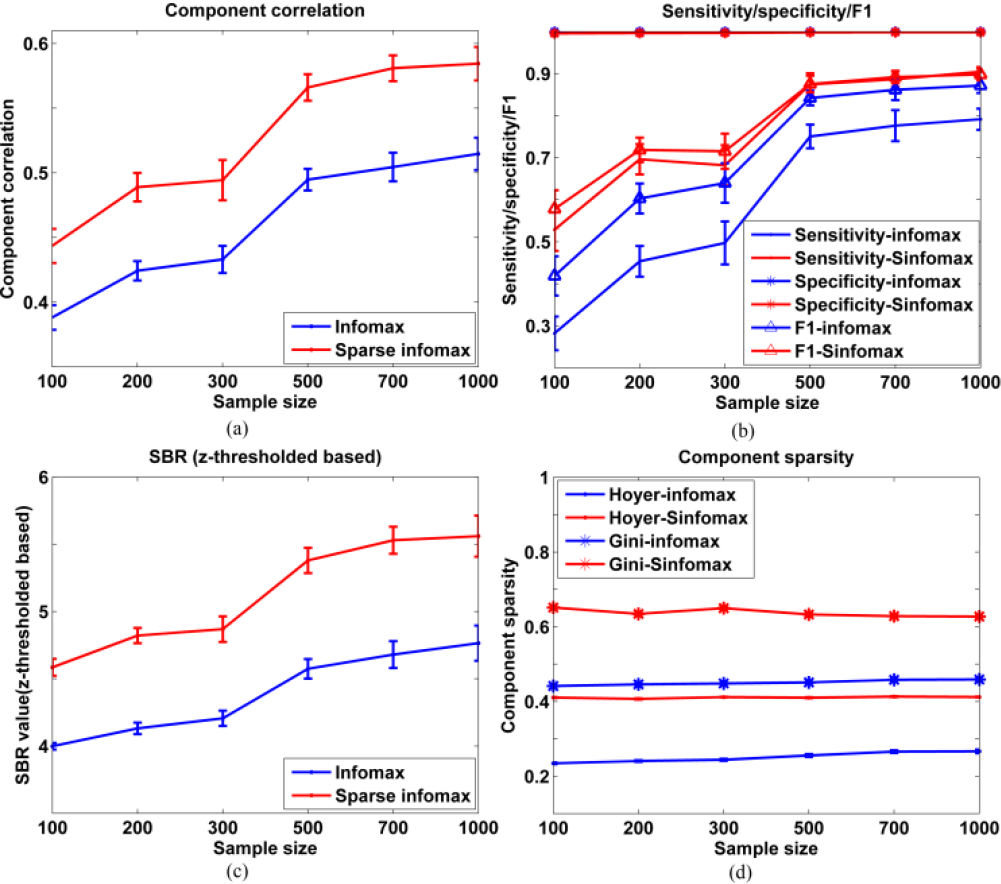
Infomax and sparse infomax performances on SNP data when varying sample size: (a) component correlation, (b) sensitivity/specificity/F1, (c) SBR (z-threshold based, |*Z*| > 3.5), (d) sparsity measures (Gini/Hoyer indices).

#### Scenario 2

sample size = 200, effect size = 3.5, number of risk SNPs varying from 50 to 200 with a step size of 50. Infomax and sparse infomax analyses were performed in the same way as in scenario 1, and the same performance measures were reported in Fig. 4.

**Fig. 4.**
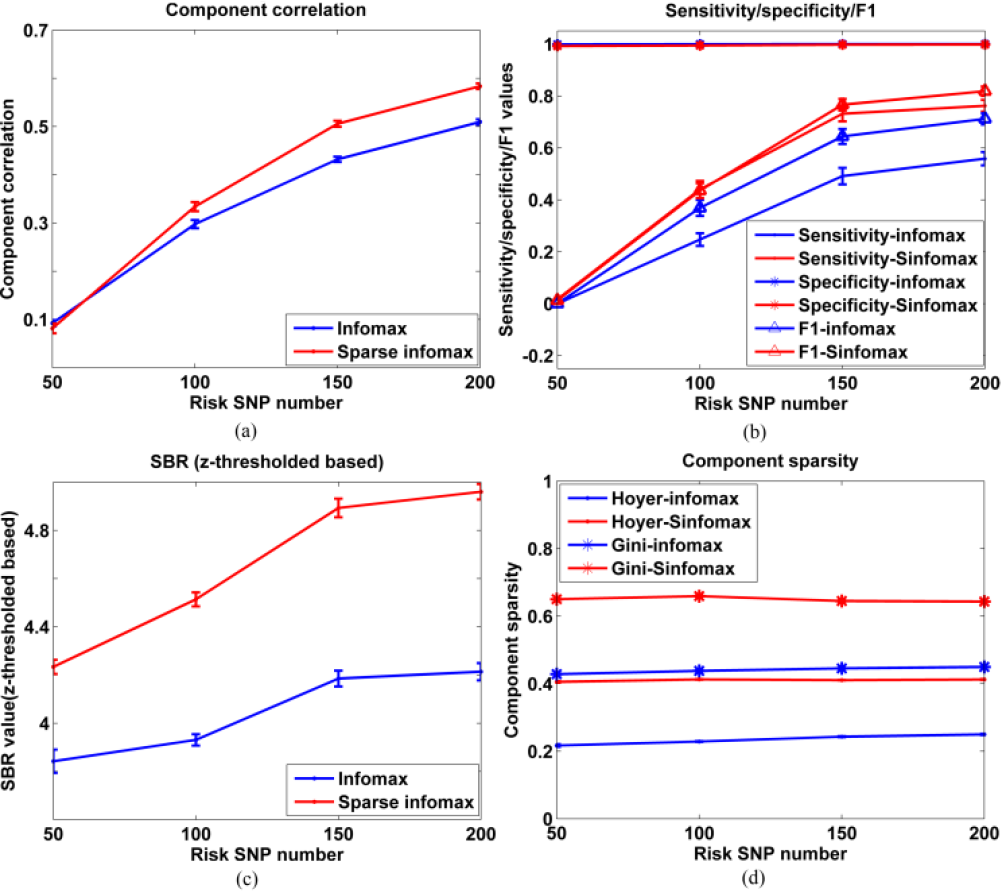
Infomax and sparse infomax performances when varying the number of risk SNPs: (a) component correlation, (b) sensitivity/specificity/F1, (c) SBR (z-threshold based, |*Z*| > 3.5), (d) sparsity measures (Gini/Hoyer indices).

#### Scenario 3

number of subjects = 200, number of risk SNPs = 150, effect size varying from 2 to 5 with a step size of 1. Same infomax and sparse infomax analyses as in scenario 1 were performed, and the same performance measures were reported in Fig. 5.

**Fig. 5.**
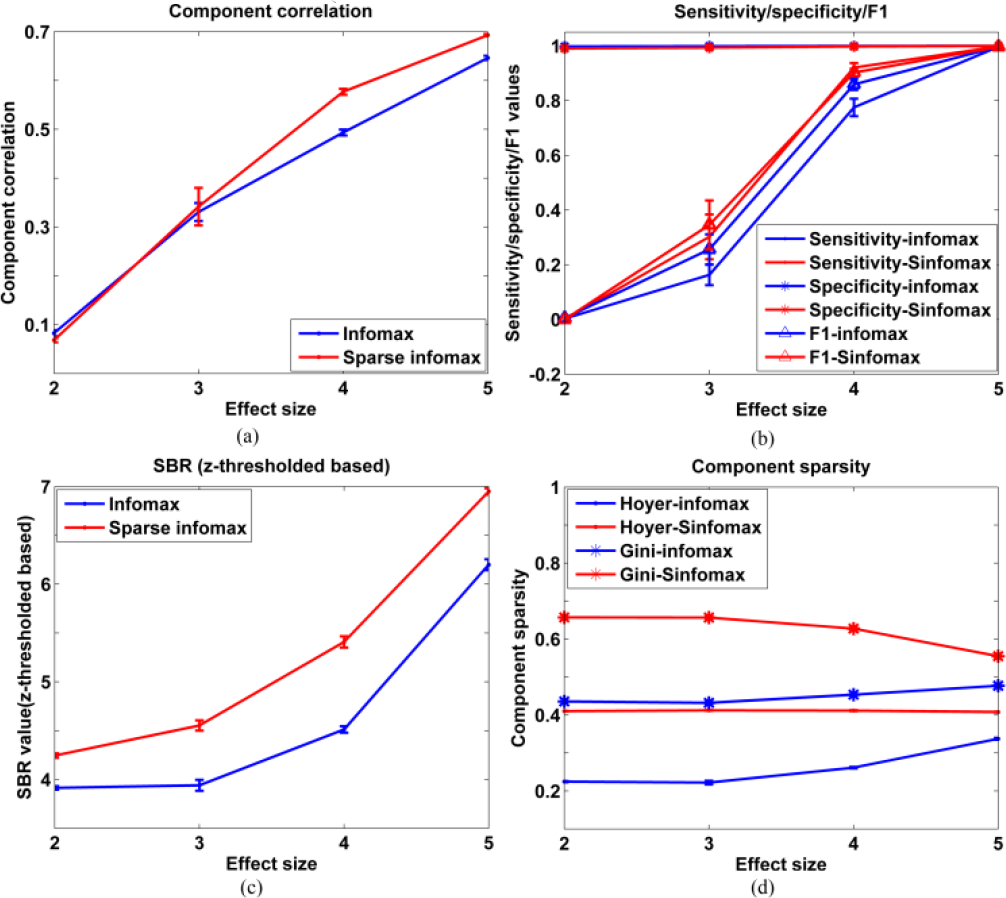
Infomax and sparse infomax performances when varying effect size (a) component correlation, (b) sensitivity/specificity/F1, (c) SBR (z-threshold based, |*Z*| > 3.5), (d) sparsity measures.

Fig. 3 shows an increasing trend in accuracy with increasing sample size, as reflected by component correlation, sensitivity/F1 and SBR values. Sparse infomax always outperforms infomax, which suggests that sparse infomax provides improved pattern detection capability over infomax. In addition, sparse infomax is tunable and, thus, can produce even sparser components. The convergence speeds are comparable between infomax and sparse infomax when varying subject number (similarly for scenarios 2 and 3).

Fig. 4 also shows an increasing trend in terms of component correlation, sensitivity/F1, and SBR as the number of risk SNPs increases. When the number of risk SNPs is 50 and 100, both infomax and sparse infomax fail to recover the true sources, which is likely caused by the fact that the PCA step removes too much variability from the data. In cases where the number of risk SNPs are larger than 100, sparse infomax has higher component correlation and sensitivity/F1 than infomax. For all cases, components estimated by sparse infomax always have higher SBR and are much sparser than those from infomax.

From Fig. 5, we can observe that, as the effect size increases, the component correlation, sensitivity/F1 and SBR increases. When effect size is low (e.g., 2 and 3), both infomax and sparse infomax fail to detect the underlying components. In the cases where effect sizes are greater than or equal to 4, sparse infomax outperforms infomax in terms of component correlation, sensitivity/F1 (which are comparable to those from infomax when the effect size is 5), SBR and sparsity measures.

## 4. DISCUSSION

A sparse infomax algorithm based on the Hoyer projection is proposed in this paper. Sparse infomax leverages the sparsity nature of sources to help suppress noise (background) and enhance signal, through a nonlinear projection, thus, expanding the solution search space as compared to traditional linear transformation solvers based on gradient optimization, and then use Tychonov-regularized least squares, taking noise into account to estimate the mixing matrix from the estimated sources. Simulation results show that recovering the independent sources with sparsity optimization can increase the component detection power for situations where the source SBR is low. This method is particularly beneficial to SNP data. The setting of the Hoyer index preset value can be subjective to data type. By default, we adaptively adjust it to make sure that components achieve the minimum SBR (e.g. 5). The minimum SBR is set according to the observations from both simulation and real data. Also, users may opt to set a higher Hoyer index preset value in order to produce sparser components.

## 5. ACKNOWLEDGE

This study was supported by the National Institutes of Health grants R01MH106655, R01EB005846, P20GM103472, P30GM122734 and NSF grant 1539067.

## REFERENCES

[1] A. J. Bell and T. J. Sejnowski, “An Information Maximization Approach to Blind Separation and Blind Deconvolution,” Neural Computation, vol. 7, pp. 1129–1159, Nov 1995.

[2] R. Y. Ge, Y. B. Wang, J. P. Zhang, L. Yao, H. Zhang, and Z. Y. Long, “Improved FastICA algorithm in fMRI data analysis using the sparsity property of the sources,” Journal of Neuroscience Methods, vol. 263, pp. 103–114, Apr 1 2016.

[3] N. Hurley and S. Rickard, “Comparing Measures of Sparsity,” Ieee Transactions on Information Theory, vol. 55, pp. 4723–4741, Oct 2009.

[4] P. O. Hoyer, “Non-negative matrix factorization with sparseness constraints,” Journal of Machine Learning Research, vol. 5, pp. 1457–1469, Nov 2004.

[5] G. Strang, COMPUTATIONAL SCIENCE AND ENGINEERING. Wellesley, MA: Wellesley-Cambridge Press, 2007.

[6] E. B. Erhardt, E. A. Allen, Y. H. Wei, T. Eichele, and V. D. Calhoun, “SimTB, a simulation toolbox for fMRI data under a model of spatiotemporal separability,” Neuroimage, vol. 59, pp. 4160–4167, Feb 15 2012.

